# TaxOnTree: a tool that generates trees annotated with taxonomic information

**DOI:** 10.1101/2020.12.24.424364

**Authors:** Tetsu Sakamoto, J. Miguel Ortega

## Abstract

Phylogenetic analysis is a widely used approach for analyzing and illustrating gene/protein/species evolution and is being benefited by the increasing number of species with their DNA/genome sequenced. Generating a phylogenetic tree with sequences from hundreds of species can be considered a routine task. However, tree visualization has been challenged to organize and bring by accessible means relevant information, e.g. taxonomy, about the sampled genes/proteins. Here we present TaxOnTree, a computational tool that incorporates and allows a quick accession of the taxonomic information of samples in a phylogenetic tree. TaxOnTree takes as input a single phylogenetic tree in Newick format containing gene/protein identifiers from NCBI or Uniprot databases in their leaves but TaxOnTree also allows users to have as input a protein identifier, a single protein in FASTA, a list of protein accessions, or a(n) (un)aligned multi-FASTA file. Non-tree inputs are submitted to a phylogenetic reconstruction pipeline implemented into TaxOnTree. The tree provided by the user or generated by the pipeline is converted to Nexus format and then automatically annotated with the taxonomic information of each sample comprising the tree. The taxonomic information is retrieved by web requests from NCBI or Uniprot servers or from a local MySQL database and annotated as tags in the tree nodes. The final tree archive is in Nexus format and should be opened with FigTree software which allows visual inspection, by branch coloring or tip/node labeling, of the taxonomic information incorporated in the tree. TaxOnTree provides prompt inspection of the taxonomic distribution of orthologs and paralogs. It can be used for manual curation of taxonomic/phylogenetic scenarios and coupled to any tool that links homologous sequences to a seed sequence. TaxOnTree provides computational support to help users to inspect phylogenetic trees with a taxonomic view, even without being taxonomy experts. TaxOnTree is available at http://bioinfo.icb.ufmg.br/taxontree.

## Introduction

Analyzing and illustrating gene/protein evolution are common tasks in research seeking for some features in a gene/protein of interest. One widely used approach is the phylogenetic analysis, which possesses robust methods for inferring the evolutionary history of genes/species. From a phylogenetic tree, one could understand the formation of the group of homologs, inspect gene duplication/deletion events or even detect non-neutral evolutionary events like positive/negative selection, coevolution, and lateral gene transfer. Although the building of a cluster of homologs is a very hard and disputed subject [1, 2], a simple similarity search (e.g. conducted by BLAST) coupled with the reconstruction of a phylogenetic tree supports a careful manual analysis of the evolutionary scenario.

In an era in which it is possible to build a phylogenetic tree with sequences from hundreds of species [3], several tools applied on the main steps of the pipeline for phylogenetic inference (e.g. sequence alignment, tree inference, and visualization) were developed or updated to support increasingly voluminous analyzes. In tree visualization, the main issue is in analyzing visually a large phylogenetic tree [4], coupled with the necessity for a rapid approach to access information about the sampled genes/proteins, such as their associated molecular function, family classification, and user metadata. During the tree analysis, taxonomic classification is one of the most consulted data since knowing the evolutionary scenario of the species in the tree is essential to identify some events of the gene evolution, such as duplication, gain, loss, and horizontal transfer, and non-neutral evolutionary process. Thus, it would be helpful if along with the diverging scenario presented by a phylogenetic tree, one could inspect the taxonomic information of the sampled proteins directly from the tree. Several bioinformatics tools were developed to aid in the visual task and provide a variety of approaches to graphically display other data along the tree [5–14]. However, currently available tools do not provide an automatic means to annotate a phylogenetic tree with taxonomic data. Here we describe TaxOnTree, a tool that adds taxonomic classification on top of a phylogenetic tree.

### Implementation

The main purpose of TaxOnTree is to add taxonomic information into a phylogenetic tree. However, users can also run TaxOnTree as a phylogenetic pipeline that uses well-established third party software for putative orthologs retrieval, sequence alignment, alignment quality analysis, and phylogenetic tree reconstruction (Figure 1). This allows TaxOnTree users to deal with several input formats that can be from a single protein accession to a tree file in Newick format. TaxOnTree source code (available at http://github.com/biodados/taxontree) was written in Perl and uses phylogenetic modules from BioPerl [15] to read and edit the tree. It can be run on a UNIX platform, but we also made TaxOnTree available as a web tool that can be accessed at http://bioinfo.icb.ufmg.br/taxontree.

**Fig 1.**
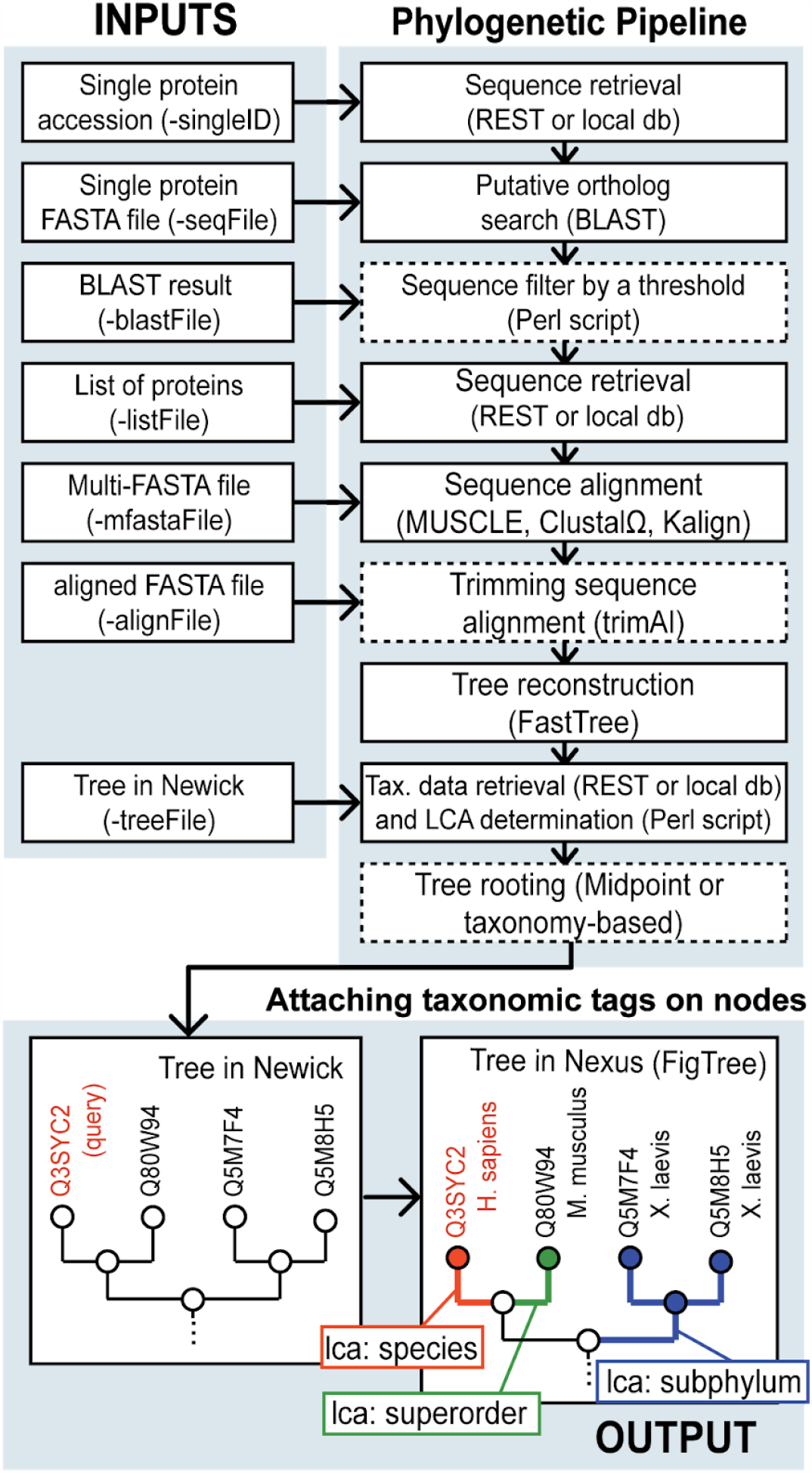
TaxOnTree workflow. Parameters used by TaxOnTree to receive each type of input are in parentheses. In the phylogenetic pipeline, dashed boxes are optional steps, and methods or software used to accomplish each step are in parentheses.

### Workflow

TaxOnTree follows the workflow schematized in Figure 1. It can receive several types of input which begin the phylogenetic pipeline at different points. To describe the whole pipeline, we will describe the operations performed by the pipeline when a single protein accession is used as input since it goes through all pipeline steps. By receiving a single protein identifier from NCBI RefSeq [16] (GI or accession number) or UniProtKB [17] (accession number or entry name) databases, our implementation retrieves its sequence by web request or from a local database and runs a BLAST search [18] to retrieve putative homologs of the query sequence. In the web tool, we provide preformatted databases from (i) UniProtKB, restricted to sequences from proteomes of reference organisms, and from NCBI RefSeq, that contains (ii) only sequences with status “Complete” to provide a more reliable comparison between sequences, or (iii) the full RefSeq database, for inspection of the whole sampling scenario. Command-line users can also build and use their own sequence databases. Protein accessions retrieved by BLAST search can be further filtered according to an identity threshold. These protein accessions, then, have their sequences retrieved and follow the next step, which is the multiple sequence alignment. Commonly used alignment software like MUSCLE [19], Clustal Omega [20], and Kalign [21] are included in the TaxOnTree package. Sequence alignment trimming is optionally performed by trimAl [22] and, then, a phylogenetic tree is generated by FastTree [23] in Newick format. Next, TaxOnTree retrieves the taxonomic information of all taxa involved in the analysis from two databases: (i) NCBI Taxonomy, the primary source of taxonomic information of sequences from GenBank; and (ii) Taxallnomy [24], a balanced taxonomic tree constructed based on the hierarchical structure provided by NCBI Taxonomy, which provides taxonomic lineages with all taxonomic ranks for all organisms. Taxonomic lineages retrieved from NCBI Taxonomy are used to determine the Lowest Common Ancestor (LCA) between the query species and each subject species in the tree. LCA here represents the most recent taxon in which two species share the same ancestor (e.g. LCA between human and mouse is the superorder Euarchonthoglires). On the other hand, the taxonomic lineages retrieved from Taxallnomy are used to avoid the appearance of missing data when querying for a taxonomic rank since taxonomic lineages provided by NCBI Taxonomy do not contain necessarily all taxonomic ranks (e.g. taxonomic lineage of Homo sapiens misses for subclass rank). At last, the tree is converted to Nexus format and all taxonomic information is stored as tags in each node of the tree. The final archive, a tree file in Nexus format, is configured to be opened in FigTree software [25]. Figtree is made freely available by authors as a Java program with versions for several OS systems and provides additional tools for tree edition and publication-ready images. The phylogenetic reconstruction pipeline implemented on TaxOnTree is restricted only to protein data. However, TaxOnTree can retrieve taxonomic information from non-protein accessions. This allows TaxOnTree to also incorporate taxonomic information in trees with nucleotide or genomic accessions in their leaves.

### Data retrieval

Data required by TaxOnTree for tree reconstruction and annotation include the taxonomic, gene, and sequence information of each protein accession in the analysis. These data are retrieved by HTTP request from NCBI [26] and/or UniProt servers. This method grants users to work with an updated data source and discards the requirement of storing a large database in a local machine. On the other hand, users can also configure and install a local database to run TaxOnTree offline. These procedures will speed up data retrieval and are appropriate for users with large demands. All necessary files to work with TaxOnTree in offline mode are hosted on our Sourceforge page (https://sourceforge.net/projects/taxontree/) and include: (i) a dump file to be loaded in a MySQL database and (ii) BLAST-formatted sequence databases. In offline mode, the internet connection will only be requested during a run if some information of a protein accession is not available in the local database.

## Results

### Inspecting taxonomic classification and relationship among species comprising the tree

After running TaxOnTree by using one of the input formats allowed, the final archive generated by the program is a tree in Nexus format with taxonomic information attached to its nodes and configured to be opened in the FigTree [25] software. To demonstrate the functionalities of those trees on FigTree, we took the Nexus file generated after submitting the human Alanine--glyoxylate aminotransferase 2 (AGXT2) accession number (Entry: Q9BYV1) to TaxOnTree. AGXT2 is a mitochondrial gene that is found in a single copy in the organisms. For this tree, the phylogenetic pipeline was configured to retrieve putative orthologs by a BLAST search against the sequence database containing proteins from Uniprot Reference Proteomes. Also, MUSCLE, trimAl, and FastTree were selected in the pipeline to respectively perform the sequence alignment, alignment trimming, and phylogenetic reconstruction. The generated Nexus file was then opened in FigTree [25] (Figure 2).

**Fig 2.**
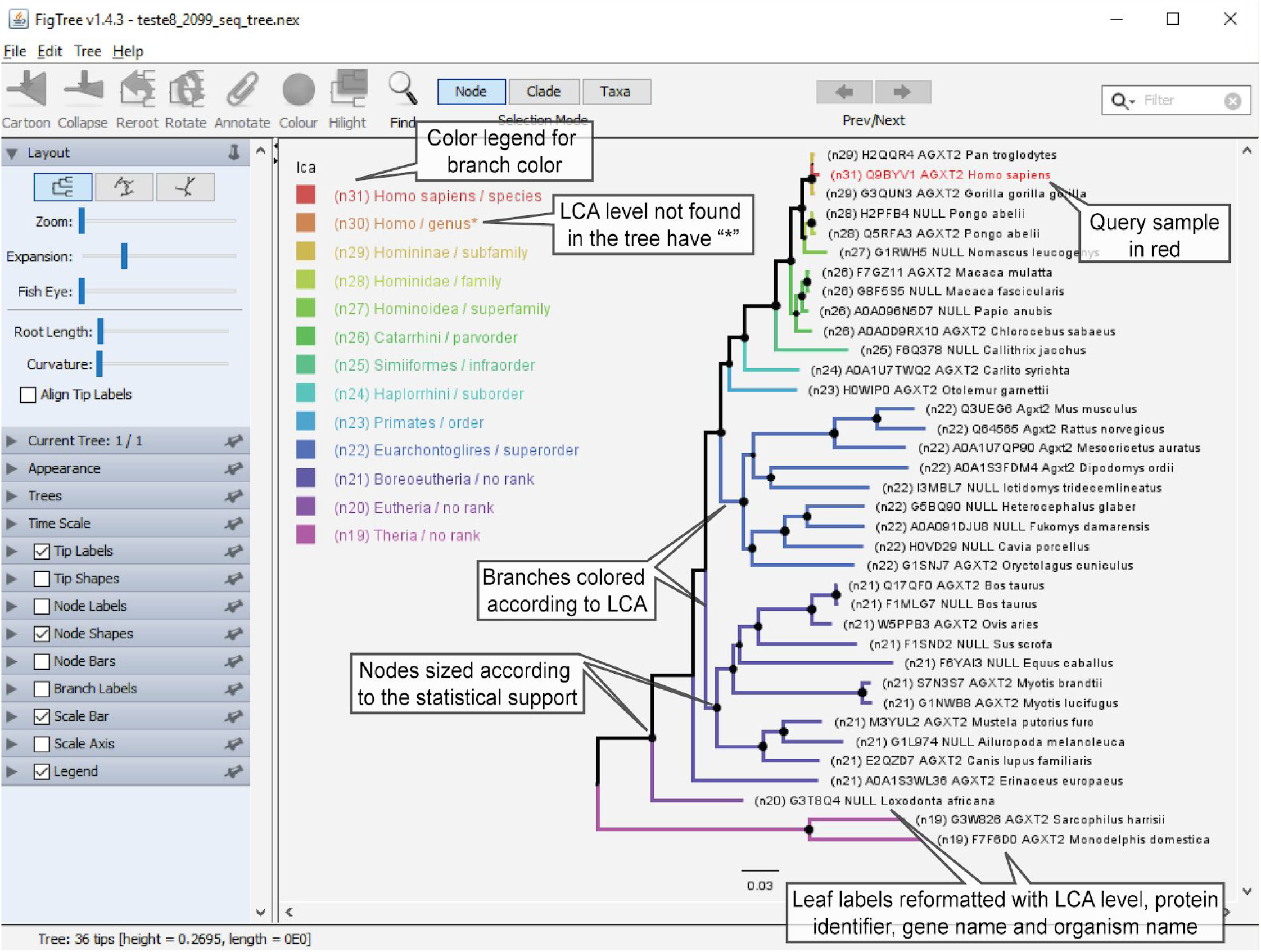
TaxOnTree output on FigTree. The Nexus file generated by TaxOnTree is formatted to be opened in FigTree. The tree was generated using the human AGTX2 (Entry Q9BYV1) as the query against the Uniprot Reference Proteome database.

In the tree displayed by FigTree, the query sample used for the BLAST search is highlighted with a red font (Q9BYV1). The tree branches are colored according to the LCA between the species of query sample (Homo sapiens) and the other species comprising the tree. The color legend refers to the branch colors and indicates the LCA level, the taxon name, and its rank. Some items in the legend may have an asterisk (*) in the end, indicating that there are no organisms in the tree which share those LCA levels with the query organism. Moreover, all nodes are sized proportionally to the statistical support of the branches. Finally, the tip labels were formatted to include some taxonomic information. By default, it displays (i) the LCA level shared with the query organism, (ii) protein accession, (iii) gene name (it displays NULL if the accession does not have a gene name), and (iv) the organism scientific name. Data in the tip label can be customized before the submission of any input to TaxOnTree.

The coloring scheme by LCA allows users to visually inspect the taxonomic relationship between the query species (in this case, *Homo sapiens*) and each of the subject species (other mammalians). Although some could be familiar with mammalian taxonomy, probably many researchers would not know at first glance that Asian monkey (*Macaca mulatta*) and mouse (*Mus musculus*) share respectively the same Parvorder and Superorder with the human. Without TaxOnTree, visual inspection of the tree would require deep knowledge of the organism’s classification.

Another coloring scheme that TaxOnTree provides is according to a taxonomic rank (Figure 3). Users can choose one of 17 taxonomic ranks (class, order, family, etc.) and visually inspect the taxonomic diversity of the selected rank in the tree. In Figure 3, the same tree in Figure 2 is displayed with branches colored according to the superorder (left tree) and order (right tree) ranks. By this, we could rapidly inspect that the samples comprising the tree are distributed into five and twelve distinct superorders and orders, respectively. Note that some superorder names begin with an abbreviation of their rank, e.g. “SpOrd_of_Didelphimorphia”. They are, actually, taxa that were created by the Taxallnomy algorithm [24], since the species that belong to these superorders do not have a taxon of superorder rank in their taxonomic lineage.

**Fig 3.**
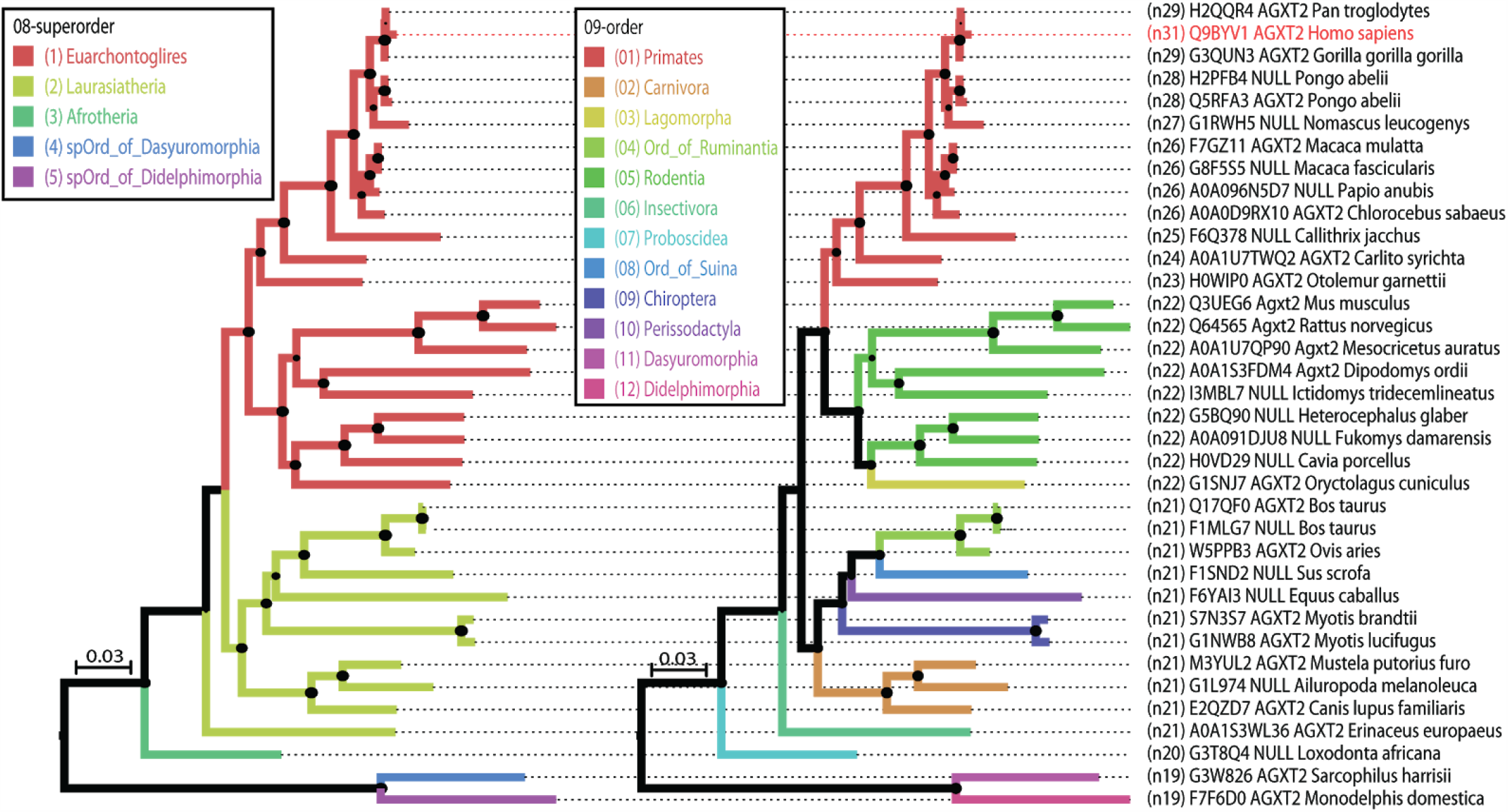
Branch coloring by a taxonomic rank. Both trees are the same gene tree of Figure 2. but their branches were colored according to the superorder (left) and order (right) ranks.

### Inspecting duplication/deletion events

An evolutionary scenario that can also be better depicted by TaxOnTree is the gene duplication/deletion events. To illustrate this, we analyzed the phylogeny of some members of the solute carrier organic anion transporter (SLCO) family using the human SLCO member 1B7 (SLCO1B7; accession: NP_001009562) as query and NCBI RefSeq as the sequence database for putative orthologs retrieval. In the resultant tree (Figure 4A). Although almost all species have the gene in a single copy, by setting the node shape on FigTree to highlight the duplication nodes, we could verify the occurrence of duplication events in clades comprising species of order Macroscelidea, “Ord of Tylopoda”, Rodentia and Primates. In Primates clade (Figure 4B), we could depict four duplication events. Two of them probably have occurred specifically in the lineages of *Carlito syrichta* and of *Cebus capucinus imitator*. The other two duplication events have occurred in the ancestor of all Simiiformes and gave rise to two more SLCO1B subfamily members, now referred to as SLCO1B1, SLCO1B3, and SLCO1B7. The inspection of deletion events in the tree is also facilitated through branch coloring. By making the branches colored according to subfamily rank, we could verify that the clade comprising the SLCO1B7 gene is lacking some branches with colors that are found in the other two SLCO1B gene clades. If the genomes of the sampled species are well-annotated, one could suggest that they were deleted in those species. But, since not all Simiiformes species have a high-quality genome sequence available, there is a chance of being a miss annotation problem. Despite this, we could easily observe that there is a good chance that the SLCO1B7 gene has been deleted in the lineage of the Colobinae subfamily since all sampled species of this subfamily lack this gene.

**Fig 4.**
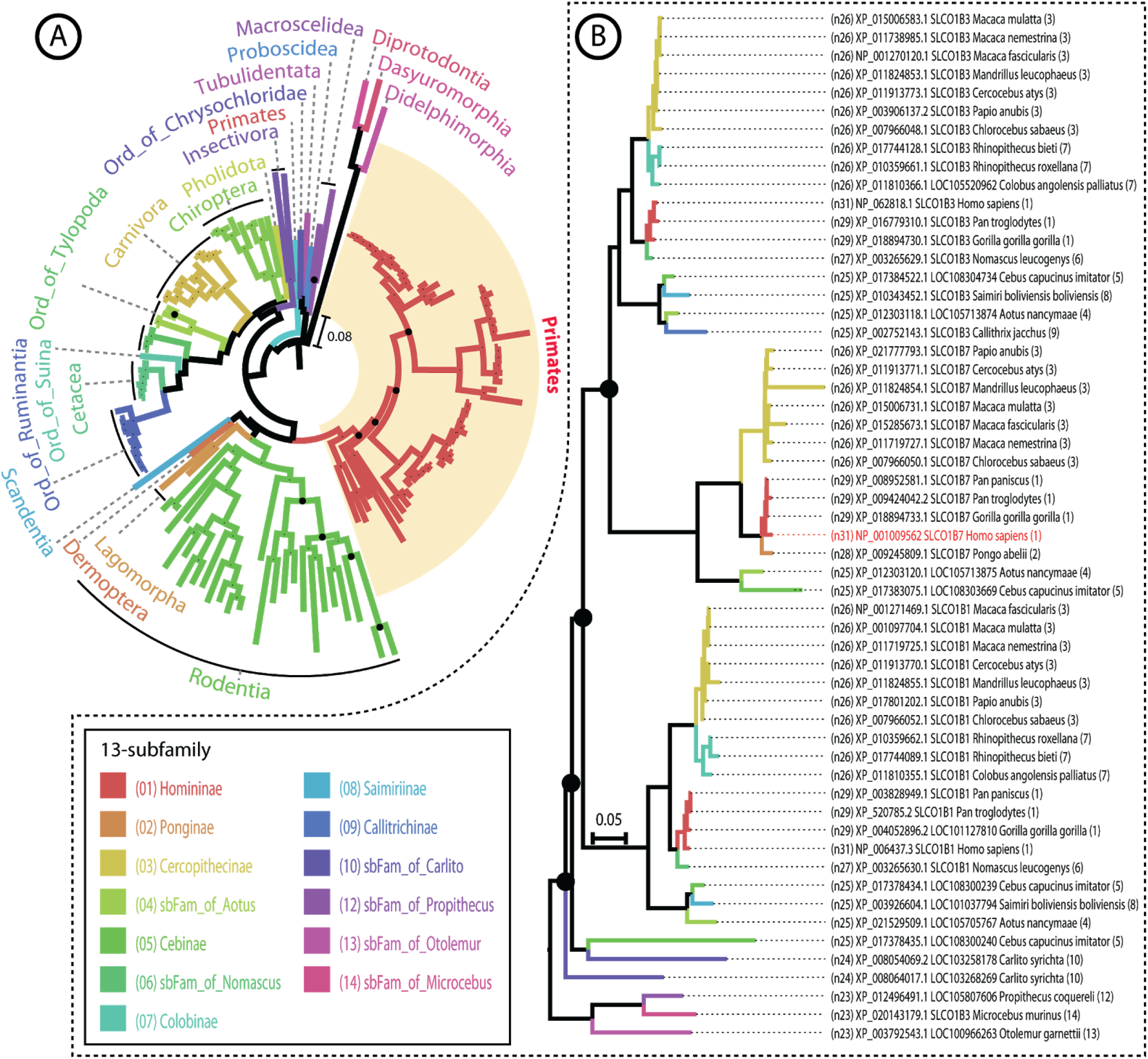
Depicting duplication/deletion events in SLCO1B protein subfamily. All highlighted internal nodes represent duplication events. (A) Full-view of the tree generated by TaxOnTree using the human SLCO1B7 (accession. NP_001009562) and RefSeq as query and sequence database, respectively. The branches are colored according to the different orders compnsing the tree. (B) Subtree detailing the branch that gathers all samples from Primates’ order. The branches are colored according to the different subfamily in the tree. The leaf names were formatted by TaxOnTree and are showing the LCA level, protein accession, gene name, species name and taxonomy code for the subfamily rank (the number found in the legend).

### Other features in TaxOnTree

### Sample filtering

Although the increasing number of samples benefits the tree inference, large trees are difficult either to analyze or to visualize. Thus, we implemented on TaxOnTree some options that allow users to filter the samples in the final tree. One of them is a filter that removes protein isoforms. Since TaxOnTree retrieves gene information from all protein accessions in the analysis, users can optionally allow TaxOnTree to keep a single isoform for each gene for further procedures. The remained protein accessions, depending on the input type, could represent the isoform of the highest similarity with the query protein (if there is a BLAST result available or if it was provided a tree file), or the first isoform found in a set of accessions or sequences provided by the user.

Other filters implemented in the TaxOnTree are based on taxonomic information. These filters can be useful in dealing with organisms that are over-represented in the database (i.e. Escherichia coli has more than 9000 genomes entries deposited in GenBank) or in selecting organisms of interest to comprise the final tree. Currently, TaxOnTree can use taxonomic information to filter the samples by:

- Limiting the number of organisms in taxonomic groups – In this filter, the user can define the number of organisms to be displayed in the taxonomic groups which comprise a category. The category can be a taxonomic rank or the LCAs shared with the query organism. For instance, if “2” and “Kingdom” are provided, TaxOnTree will display up to two organisms on each Kingdom comprising the tree.
- Establishing an LCA level range – In this filter, TaxOnTree will display only those samples in which the level of the LCA shared with the query organism is in the range defined by the user. The user can opt to display those organisms with LCA level above or below the threshold or combine both parameters to display samples from a range of LCA level.
- Listing a set of organisms to be displayed in the tree – Users can provide a list of TaxonomyID to display only those samples that belong to organisms that have their TaxonomyID listed.

### Leaf name customization

Leaves are the primary source for accessing data of a specified sample. If their names contain appropriate information, they largely help users on the tree interpretation without consulting other data sources. However, leaf names are generally displayed as the FASTA header of the alignment file and mostly contain only the protein accession number. Users are also confronted by restrictions of some well-known phylogenetic reconstruction software in naming samples, such as the character number limit or the constraint of some special characters. TaxOnTree meets this issue by allowing users to customize the information to be displayed in the leaves of a tree before the job submission. Data that can be added to the leaf name by TaxOnTree includes:

- Primary ID – Accession number (for NCBI accession) or Accession (for Uniprot accession);
- Alternative ID – GI number (for NCBI accession) or Entry
- Name (for Uniprot accession);
- Fasta header – Original name extracted from FASTA file;
- LCA level – Corresponds to the number of taxonomic nodes found from the root to the LCA shared by the query and the subject organisms. The higher this value is, the closer is the taxonomic relationship between both organisms.
- Gene name – Official gene symbol provided by HGNC;
- Species name – Scientific name of the species or strain;
- Taxonomy rank name – Displays the taxon name of the selected taxonomic rank.
- Taxonomy rank code – Displays the code assigned to the taxon of the selected taxonomic rank. The code is a number assigned to each taxon of a taxonomic rank which reflects the similarity of those taxa with the query sample. The code ranges from 1 (taxon containing the query organism) to the number of taxa in the selected rank. The code is also displayed in the legend of the trees in which have taxa of a taxonomic rank highlighted;

Alternatively, users can also set FigTree to display information stored on tags of each leaf node on the “Tip labels” option, which could be one of the 17 taxonomic ranks, the information about LCA level, or other data provided by the user (see Figures 5C and 5D).

**Fig 5.**
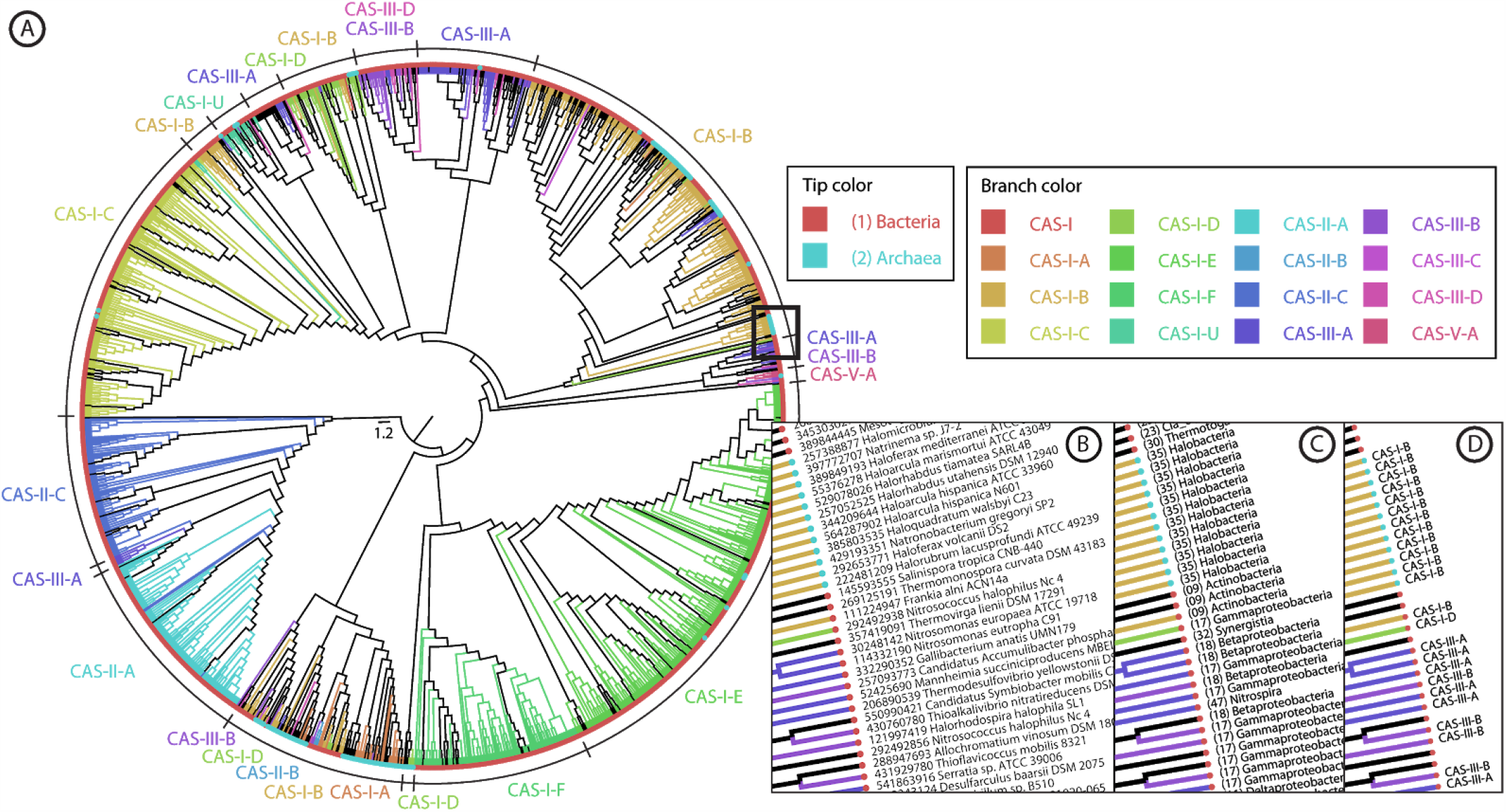
Incorporating additional data to the tree. (A) Phylogenetic tree of the Cast gene from CRISPR-Cas loci depicted in [27] and processed by TaxOnTree. Branches and tips are colored according to, respectively, the CRISPR-Cas group and the Superkingdom that each sample belongs to. The thick rectangle delimits the area of the tree which is shown In more details in (B), (C) and (D). The tip labels were configured to show the GI and the scientific name of the organism that the sample belongs to (B). but during the tree inspection in FigTree, users can configure the information shown on the tips to make it display data stored on the tags. This could be either (C) taxonomic data embedded by TaxOnTree (the lips are showing the Class rank of each sample) or (D) the data provided by the user (in this case, the CRISPR-Cas classification). Samples In the tree without a CRISPR-Cas classification are those In which their classification Is unknown, ambiguous, or partial. The tree file and the classification of each sample In the tree are made available as supplementary files in [27].

### Incorporating user annotation/metadata to the tree

Besides taxonomic data, TaxOnTree also allows users to include other annotations and view them in the tree along with the taxonomic information. For this, users only have to provide a tab-delimited file containing the names of the samples comprising the tree in the first column and the information associated with them in the second one. To illustrate this, we resorted to a published work that performed a phylogenetic analysis of the Cas1 gene, a component of *CRISPR-Cas* loci [27]. In this work, the tree was generated to verify if the new *CRISPR-Cas* loci classification system proposed by the authors (based on the architecture of *cas* loci) is in agreement with the phylogeny of the Cas1 gene. By submitting the Cas1 gene tree to TaxOnTree along with the tab-delimited file containing the CRISPR-Cas classification of each sample, we generated a Nexus file containing CRISPR-Cas classification data on their nodes, along with the taxonomic data. Therefore, by opening the Nexus file on FigTree we could not only reproduce the tree image displayed in the referred article but also provide a tree visualization that allows rapid access to both taxonomic and user-provided data (Figure 5).

### Tree rooting

Most of the software available for phylogenetic reconstruction generates an unrooted tree. Thus, the classical midpoint rooting method is implemented in TaxOnTree and used as the default method for tree rooting. However, if it is known the taxonomic background of the samples in the tree, one could manually inspect the tree to find a more appropriate root point. Since TaxOnTree retrieves the taxonomic information of the samples comprising the tree, we implemented an algorithm that takes advantage of taxonomic data to automatically look for a suited branch and root it. In this method, denominated as taxonomy-based rooting method (Figure 6), each pair of organisms in the tree has their taxonomic relationship measured by the LCA level. Using this measure, the method visits each internal node and identifies which of the branches leaving the node are more suited to be an ancestor branch. A branch is considered more suited to be a putative ancestor if it forms a cluster in which the LCA level between the species in the cluster is the highest one, compared to the LCA level of other clusters formed if another branch were considered to be the ancestor. Then, the algorithm evaluates again each internal node by counting how many putative ancestor branches will be in fact an ancestor branch if a root node were to be created on it. The branch that resulted in the highest count (score) will have a root node created on it. It is worth to note that the current implementation of this method will attempt to find a root in the tree which brings together all samples from organisms that are distantly related to the rest of the organisms in the tree and does not correct the root position if there is an artifact in the tree, i.e. a bacterial sample next to a human sample. Therefore, we recommend this approach for those controlled experiments in which it is known a priori that there is a set of distantly related samples in the tree.

**Fig 6.**
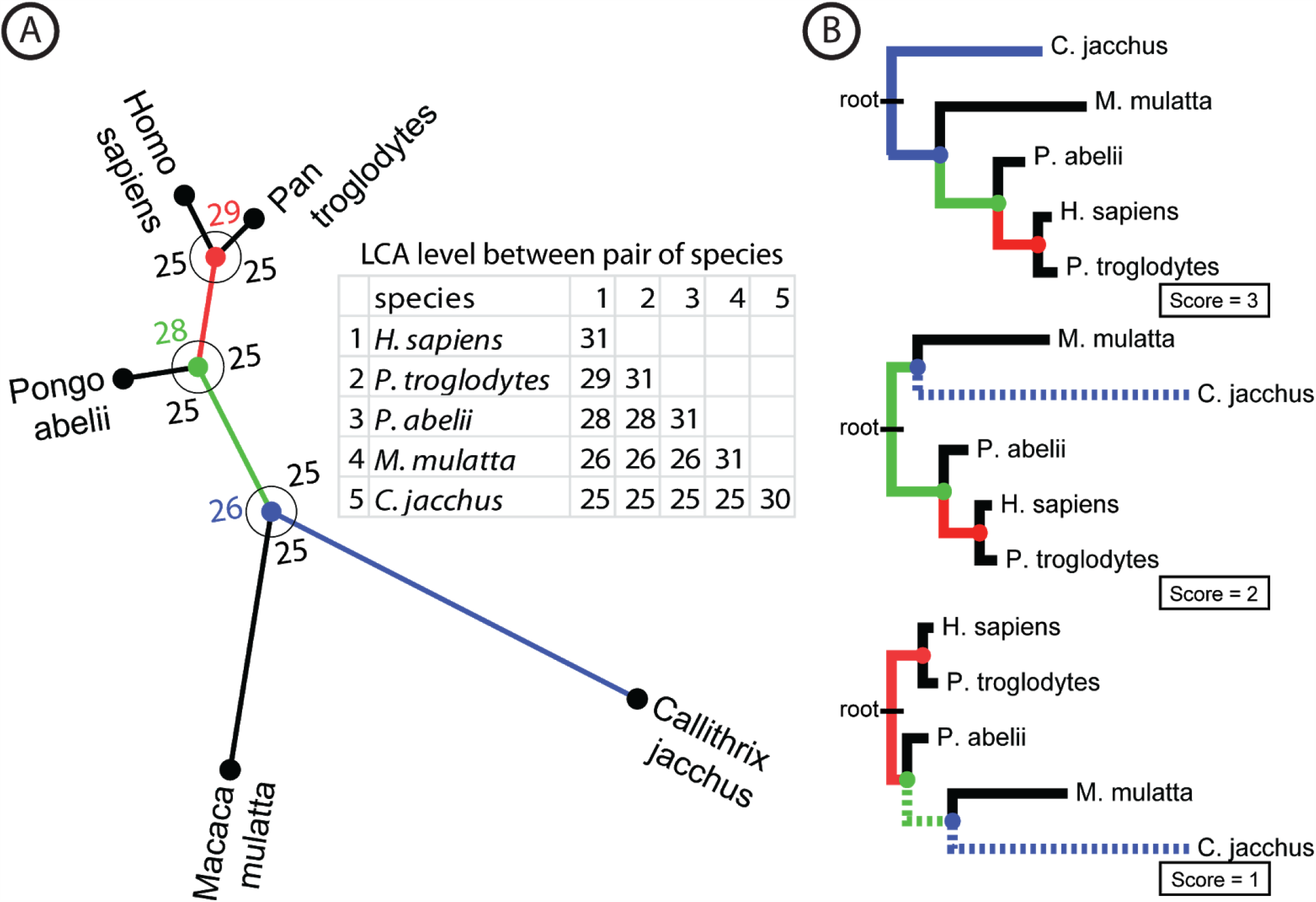
Taxonomy-based rooting method. By taking an unrooted tree (A), the algorithm firstly evaluates each internal node (identified with a color) determining which branch leaving the node is more suited to be the ancestor one based on the taxonomic relationship between species measured by LCA level (table). The numbers in the internal node on each arc indicate the LCA level of the set of organisms comprising the two branches linked by the arc. In other words, it corresponds to the LCA level of the cluster formed when the branch opposing the arc is considered as the ancestor one. The branch that forms a cluster with the highest LCA level (colored branches) is considered to be a putative ancestor branch. Then, (B) the algorithm simulates the rooting process on these branches and chooses for the final rooted tree the one with the largest amount of putative ancestor branches in agreement (the tree with score=3).

### Taxonomy report

Another output generated by TaxOnTree is a tab-delimited file containing a summary of the taxonomic clusters comprising the tree (Figure 7). In this report, the user can verify how many taxonomic clusters have formed in each taxonomic rank and obtain information about each cluster including the number of distinct species comprising it, the branch statistic supporting the cluster, and the mean of the distances between the query sequence and the leaves in the cluster. The report also allows users to have a trace about the tree topology by informing the clusters comprising the sister group and the immediate outgroup of each cluster. Users with programmatic skills could take advantage of this report to analyze numerous trees. As an example, we downloaded 14,526 maximum likelihood trees from the OrthoMam database [28] and submitted them to TaxOnTree as input. The OrthoMam has a collection of rooted phylogenetic trees comprising sequences from mammals that have their genome sequenced. All generated taxonomy report was then processed by a script that classified each taxon of superorder rank according to its cluster organization as: (i) “single”, if the taxon form a monophyletic clade; (ii) “partial in”, if the branch comprising all samples of a superorder taxon has few species from other taxa; (iii) “partial out”, if most species from this taxon cluster together; or (iv) “multiple”, if species from this taxon do not cluster together. By analyzing the four eutherian superorders (Euarchontoglires, Laurasiatheria, Afrotheria, and Xenarthra), we verified that 5,310 gene trees have all those superorders forming a unique cluster. If we consider the categories “partial in” and “partial out” in the count, the number of gene trees increases to 7,517. These results and gene trees from OrthoMam processed by TaxOnTree can be accessed at http://bioinfo.icb.ufmg.br/taxontree/table_orthomam.html.

**Fig 7.**
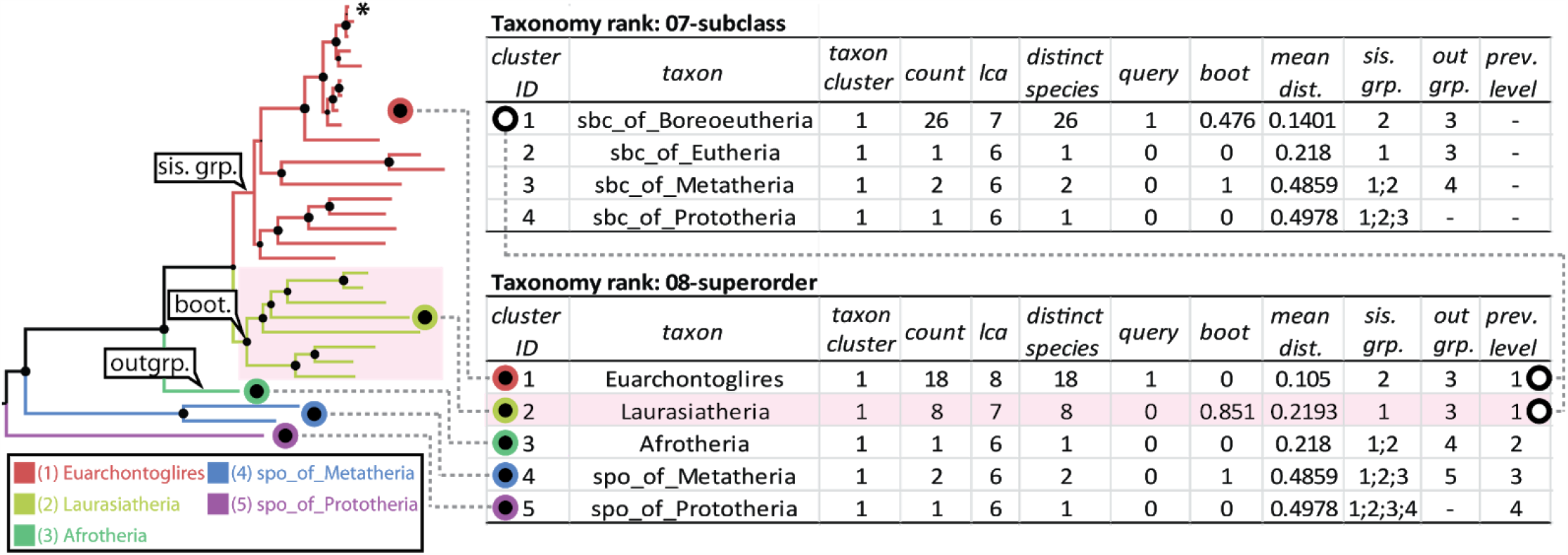
Taxonomy report output. (left) A sample phylogenetic tree colored according to the superorder rank. The position of the query sequence is indicated by an asterisk, (right) Taxonomy report generated from the sample tree. Here is displayed only the information about dusters formed in the subclass and superorder ranks, but the report can be extended to all 17 taxonomic ranks. In the report, all five dusters in the tree are reported in the correspondent rank (superorder). From the report, we could verify that the highlighted duster (Laurasiathena) forms a single cluster of 8 leaves of 8 distinct species. The LCA level, using the taxonomy lineage from Taxallnomy, the mean distance between query species and laurasiatherian species, and the statistical support (boot) for the branch leading to the duster are, respectively, 7, 0.2193, and 0.851. From the report, we could also have a general idea of the position of the duster in the tree by verifying the last three columns. The cluster Laurasiatheria has the cluster 1 (Euarchontoglires) as sister group (sis. grp.), and the cluster 3 (Afrotheria) as immediate outgroup (outgrp.). Moreover, Laurasiatheria is part of cluster 1 (prev. level) of the previous taxonomic rank level (subclass) together with Euarchontoglires.

### SVG output

Users can also generate a vector graphic output of the phylogenetic tree in SVG format. For this, we included in the TaxOnTree package a script that uses the FigTree’s [25] command-line functions that generates the tree in SVG format. This script has as input the Nexus file generated by TaxOnTree and allows the user to choose the tree coloring mode that can be by LCA or by a taxonomic rank. This output could be useful for users that may want to provide their tree graphics on the web as demonstrated in our example for large-scale tree analysis in the section “Taxonomy report”.

## Discussion and Conclusion

Automatic annotation of the phylogenetic tree with taxonomic information can be performed with some publicly available tools like PhyloView [29], iToL [8], and ETE toolkit [30]. iToL and ETE toolkit are both well-documented applications and have functions that allow automatic tree annotation by taxonomy. However, the taxonomic annotation for both applications requires users to provide the taxonomy ID associated with each leaf and/or each node in the tree. Since taxonomy ID is not always easily accessible, it implies more effort to achieve a similar result. PhyloView is a tool that most resembles TaxOnTree functionality. It is a web tool that takes a tree in Newick format as input, retrieves taxonomic information through web request by only the need of the protein accession, and colors the tree according to the taxonomic group comprising the tree. Despite those similarities, TaxOnTree provides additional features like a command-line interface for batch analysis, leaf name formatting, and tree edition which is made possible using all functionalities provided by FigTree [25]. Similarly, software for tree annotation, like ColorTree [31] and MixtureTree Annotator [32], can also provide tree visualization with the taxonomic information highlighted. However, as these applications were developed for general purpose, users will need more effort to set up the program to generate a similar tree visualization. Moreover, although TaxOnTree was developed mainly for annotating the tree with taxonomic information, it also accepts other data to be included in the tree as tags for visualization, generating a result similar to the mentioned applications, but allowing simultaneous inspection of both taxonomic and user-provided data. A feature that we have as a perspective for TaxOnTree improvement is linked to the format of the final tree generated. Currently, the annotated tree file provided by TaxOnTree is a Nexus file that is exclusively formatted to be opened in FigTree. We aim to implement new functions on TaxOnTree to generate other tree formats, like phyloXML, widening the options of a tree visualization tool in which users could use to open trees generated by TaxOnTree. TaxOnTree was developed to facilitate access to taxonomic information during a phylogenetic analysis by automating several procedures that could demand a large effort if done manually. The functionalities provided by this tool attend either those users interested in analyzing a few genes or those users with large-scale demand. The final tree file instantly gives users a taxonomic background in a phylogenetic tree, thus allowing the inspection of the species evolution as inferred by taxonomic classification along with the gene evolution.

## Availability and requirements

Project name: TaxOnTree

Project home page: http://bioinfo.icb.ufmg.br/taxontree Operating system(s): UNIX

Programming language: Perl

Other requirements: Internet connection and/or MySQL License: GNU GPL3

Any restrictions to use by non-academics: none

## Author Contributions

TS developed and designed the software and JMO supervised this project. All authors read and approved the final manuscript.

## Acknowledgments

This work has been supported by FAPEMIG through Pós-Graduação em Bioinformática ICB/UFMG, CAPES (Coordenação de Aperfeiçoamento de Pessoal de Nível Superior) and CNPq (Conselho Nacional de Desenvolvimento Científico e Tecnológico).

